# Re-annotation of SARS-CoV-2 proteins using an HHpred-based approach opens new opportunities for a better understanding of this virus

**DOI:** 10.1101/2023.06.06.543855

**Authors:** Pierre Brézellec

## Abstract

Since the publication of the genome of SARS-CoV-2 – the causative agent of COVID-19 – in January 2020, many bioinformatic tools have been applied to annotate its proteins. Although effcient methods have been used, such as the identification of protein domains stored in Pfam, most of the proteins of this virus have no detectable homologous protein domains outside the viral taxa. As it is now well established that some viral proteins share similarities with proteins of their hosts, we decided to explore the hypothesis that this lack of homologies could be, at least in part, the result of the documented loss of sensitivity of Pfam Hidden Markov Models (HMMs) when searching for domains in “divergent organisms”. In order to improve the annotation of SARS-CoV-2 proteins, we used the HHpred protein annotation tool. To avoid “false positive predictions” as much as possible, we designed a robustness procedure to evaluate the HHpred results. In total, 6 robust similarities involving 6 distinct SARS-CoV-2 proteins were detected. Of these 6 similarities, 3 are already known and well documented, and one is in agreement with recent crystallographic results. We then examined carefully the two similarities that have not yet been reported in the literature. We first show that the C-terminal part of Spike S (the protein that binds the virion to the cell membrane by interacting with the host receptor, triggering infection) has similarities with the human prominin-1/CD133; after reviewing what is known about prominin-1/CD133, we suggest that the C-terminal part of Spike S could both improve the docking of Spike S to ACE2 (the main cell entry receptor for SARS-CoV-2) and be involved in the delivery of virions to regions where ACE2 is located in cells. Secondly, we show that the SARS-CoV-2 ORF3a protein shares similarities with human G protein-coupled receptors (GPCRs), such as Lutropin-choriogonadotropic hormone receptor, primarily belonging to the “Rhodopsin family”. To further investigate these similarities, we compared Prominin 1 and Lutropin-choriogonadotropic hormone receptor to a set of viral proteins using HHPRED. Interestingly, Prominin 1 showed similarities with 6 viral Spike glycoproteins, primarily from coronaviruses. Equally interestingly, Lutropin-choriogonadotropic hormone receptor showed similarities with 23 viral G-protein coupled receptors, particularly from Herpesvirales. We conclude that the approach described here (or similar approaches) opens up new avenues of research to better understand SARS-CoV-2 and could be used to complement virus annotations, particularly for less-studied viruses.

## Introduction

A significant fraction of the proteins expressed by viruses often lack homologs. These proteins are termed “orphan” to emphasise that no homologs are detected, or “taxonomically restricted” to indicate that they have no detectable homologs outside a given taxon (Kuchibhatla *et al*., 2014). SARS-CoV-2 (Severe Acute Respiratory Syndrome Coronavirus 2), the causative agent of COVID-19, is no exception. According to UniProt (UniProt Consortium, 2021), this virus expresses 17 proteins (see Supplemental file 1 for more details). If we consider the Pfam annotations (Mistry *et al*., 2021, http://pfam-legacy.xfam.org/) of the proteins expressed by this virus, we observe that *i*/ 4 of these 17 proteins are not Pfam annotated, *ii*/ the other 13 proteins are annotated by a set of 40 domains, 39 of which are strictly associated with viruses (the Macro domain being an exception to the rule). This clearly shows that SARS-CoV-2 domains are mostly similar to viral domains (97.5% ((39/40)*100) which are generally poorly annotated.

These results can be interpreted in two different (but complementary) ways:

1. / This virus, like many viruses, essentially contains virus-like proteins that are only present in viruses and not elsewhere,
2. / As it has been established that *i*/ some viral proteins show similarities to some proteins of their host and that *ii*/ this “molecular mimicry” is increasingly recognised (Elde & Malik, 2009), this lack of homologies outside of viral taxa can also be seen, at least in part, as a consequence of weaknesses in annotation methods.

It has been shown that HMMs stored in Pfam can lack sensitivity when searching for domains in “divergent organisms” (where the relevant signals become too weak to be identified (Terrapon *et al*., 2012)). We thus decided here to explore the second way. We naturally turned to HHpred which is known to be an efficient tool for remote protein homology detection and can be easily used via a fast server (Gabler *et al*., 2020). HHpred offers many possibilities such as searching for homologs among all proteins in an organism. HHpred is based on HHsearch and HHblits, which perform pairwise comparison of HMM profiles. Given their proven efficiency, HHsearch and HHblits have been used for some years to annotate viruses, and in particular accessory proteins of coronaviruses (Forni *et al*., 2022). They have also been used to model proteins structures expressed by SARS-CoV-2 (O’Donoghue *et al*., 2021) using related 3D structures in the PDB, *i*.*e*., structures determined for other coronaviruses, such as SARS-CoV or MERS-CoV, as well as many structures from more distantly related viruses, such as those causing polio or foot-and-mouth disease. However, the two previous works limited the homology search to viral proteins. Here, using an available database of HMMs specific to *Homo sapiens* proteins, we directly searched – using HHpred - for homologs of SARS-CoV-2 proteins in human. Thus, what was previously achievable at the Pfam domain level (for instance) now extends to human proteins.

To avoid “false positive predictions” as much as possible, we designed a procedure, mainly based on two ideas suggested in (Gabler *et al*., 2020) but not implemented, to assess the robustness of HHpred results. Using HHpred and this procedure, we detected 6 robust similarities.

## Materials and Methods

### SARS-CoV-2 protein sequences

The 17 proteins studied in this article were extracted from UniProt (https://www.uniprot.org/, UniProt Consortium, 2021). UniProt provides polyproteins 1a (pp1a) and 1ab (pp1ab) as two separate entries. The pp1ab polyprotein is cleaved to form 15 shorter proteins; the first 10 proteins, *i*.*e*., NSPs 1-10, are also cleaved from pp1a; NSPs 12-16 are unique to pp1ab. The list of proteins is given below. For each protein, we give its “Recommended Name”, its “Short Name”, its “AC - Uniprot ID”, and its length:

Replicase polyprotein 1a / pp1a / P0DTC1 - R1A_SARS2 / Length 4,405

Replicase polyprotein 1ab / pp1ab / P0DTD1 - R1AB_SARS2 / Length 7,096

Envelope small membrane protein / E; sM protein / P0DTC4 - VEMP_SARS2 / Length 75

Membrane protein / M / P0DTC5 - VME1_SARS2 / Length 222

Nucleoprotein / N / P0DTC9 - NCAP_SARS2 / Length 419

Spike glycoprotein/ S glycoprotein / P0DTC2 - SPIKE_SARS2 / Length 1,273

ORF3a protein/ ORF3a / P0DTC3 - AP3A_SARS2 / Length 275

ORF3c protein / ORF3c / P0DTG1 - ORF3C_SARS2 / Length 41

ORF6 protein / ORF6 / P0DTC6 - NS6_SARS2 / Length 61

ORF7a protein / ORF7a / P0DTC7 - NS7A_SARS2 / Length 121

ORF7b protein / ORF7b / P0DTD8 - NS7B_SARS2 / Length 43

ORF8 protein / ORF8 / P0DTC8 - NS8_SARS2 / Length 121

ORF9b protein / ORF9b / P0DTD2 - ORF9B_SARS2 / Length 97

Putative ORF3b protein/ ORF3b / P0DTF1 - ORF3B_SARS2 / Length 22

Putative ORF3d protein/ _ / P0DTG0 - ORF3D_SARS2 / Length 57

Putative ORF9c protein / ORF9c / P0DTD3 - ORF9C_SARS2 / Length 73

Putative ORF10 protein / ORF10 / A0A663DJA2 - ORF10_SARS2 / Length 38

### Sequence similarity searches

For remote homology detection, we used HHpred (Gabler *et al*., 2020). First, starting from single sequences or multiple sequence alignments (MSAs), it transforms them into a query HMM; using this HMM, it then searches the Uniclust database30 and adds significantly similar sequences found to the query MSA for the next search iteration. This strategy is very effective in detecting remotely homologous sequences but, as the user guide points out (https://github.com/soedinglab/hh-suite/wiki), “the higher the number of search iterations, the greater the risk of non-homologous sequences or sequence segments entering the MSA and recruiting other sequences of the same type in subsequent iterations”. To avoid this problem, we set the number of iterations to 0, *i*.*e*. the parameter “MSA generation iterations” was set to 0. The default settings were used for the other parameters. Note that we also briefly present in the Results section the HHpred results obtained using the default setting for “MSA generation iterations”, *i*.*e*. 3 (iterations).

Finally, it is important to note here that we use HHpred to look for similarities independently of the mechanisms underlying these similarities, *i*.*e*. homologies, horizontal transfers (e.g. obtained by “recombination” between SARS-CoV-2 and its current host, between ancestors of SARS-CoV-2 and their hosts, between SARS-CoV-2 and another virus, *etc*.), convergent evolutions, *etc*.

### Procedure for assessing the robustness of HHpred results

According to (Gabler *et al*., 2020), when the reported probability value for a hit is greater than 95%, homology is highly probable. Since viral and human proteins are being compared here, it can be assumed that the 95% threshold is too high to detect similarities. In order to be more sensitive, while controlling specificity (*i*.*e*. avoiding “false positive predictions”), we have devised a procedure that we describe below. Its purpose is to assess the robustness of the results provided by HHpred. It is based on two ideas described in (Gabler *et al*., 2020) (section “Understanding Results”) but not taken into account in HHpred.

This procedure is divided into 4 steps. From an algorithmic point of view, this procedure can be described as a “gready search algorithm”. It is performed for each protein expressed by the SARS-CoV-2 virus (see Figure) (note that this was done by hand, as there were few data to process):

**Figure 1.**
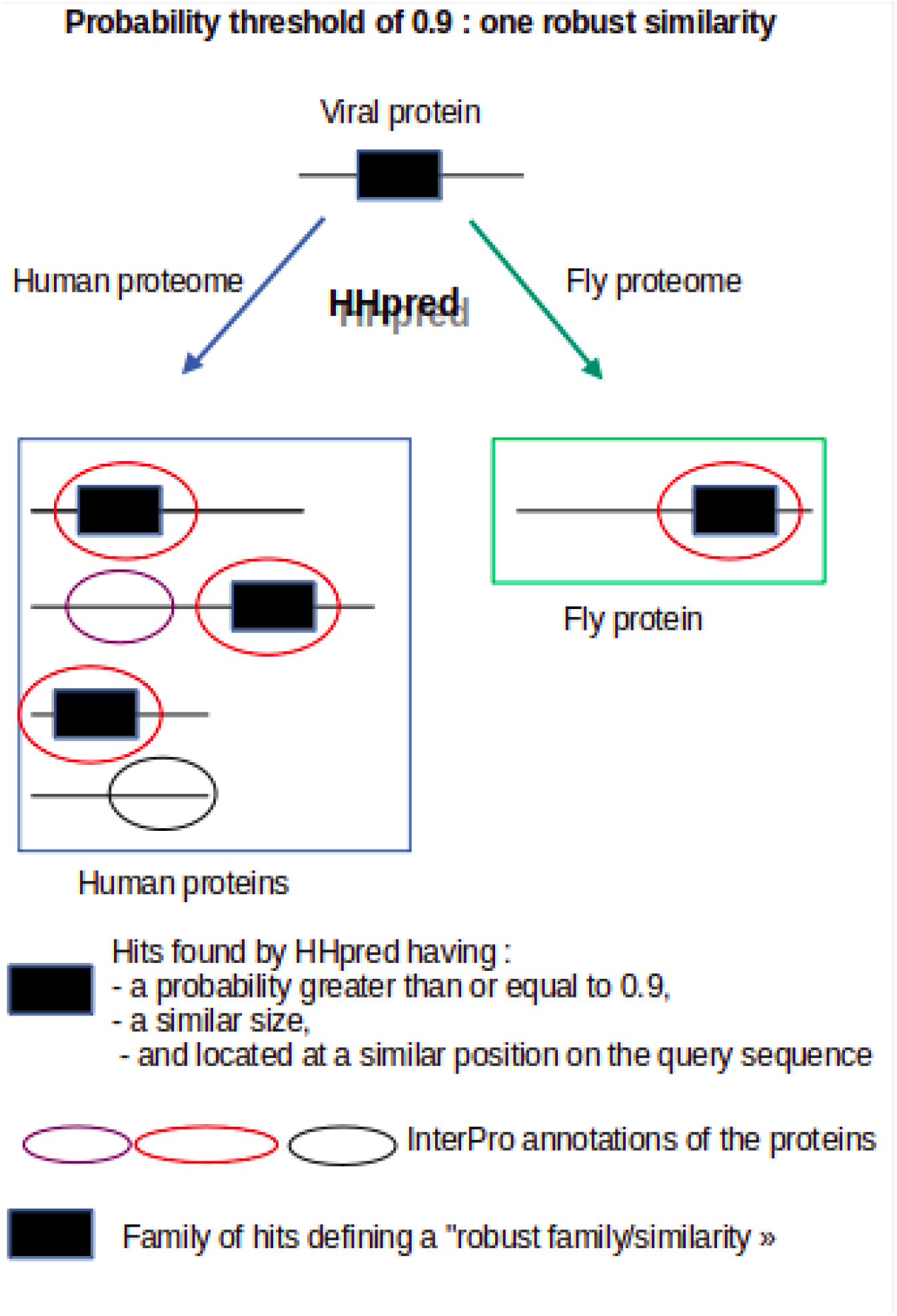
Using HHpred, a viral protein is compared to the human proteome and to a set of other proteomes, called “test” proteomes, which include the fly proteome. The probability threshold was set at 0.9, so only hits with a probability value of 0.90 or greater are considered relevant here. 4 homologous hits (*i*.*e*., hits of similar sizes and located at a similar position on the query sequence) exceeding the given threshold were found by HHpred (black boxes): 3 are found in humans and one in flies; the InterPro annotation of all the “black box” hits are the same (red oval); as the annotations of all these homologous hits are identical and at least one of these hits belongs to a test proteome, the corresponding family of homologous hits is considered to be a “robust/similar family”; this similarity will be used to annotate the corresponding hit on the viral protein.

1./ For a given SARS-CoV-2 protein, hereafter referred to as “query”, HHpred is run (using the default parameters, except for the “MSA generation iterations” parameter which we set to 0, see section above) on the *Homo sapiens* proteome of HHpred.

2.1 / The examination of the results provided by HHpred starts with the probability threshold of 0.95. Hits with a probability greater than or equal to 0.95 are selected. If no hits meet this constraint, the threshold is successively lowered to 0.9, 0.85 and finally to 0.80. As soon as a threshold satisfies the constraint (*i*.*e*. there is at least one hit with a probability greater than or equal to the threshold), all hits above the threshold are selected. If no threshold satisfies the constraint, we consider that no similarity between the query and the human proteins can be detected.

2.2./ All previously selected hits are collected in a list and ranked from highest to lowest probability. The best hit is then used as a seed to build a family of hits as follows: hits located at the same position as this best hit on the query sequence and of similar size to it (+-5 amino acids for a best hit of length < 150, and +-15 for a best hit of length > 150) feed the family under construction and are removed from the list; hits that overlap the seed are also removed from the list. The highest hit in the updated list is used as the “new” seed and the process continues until the list is empty. As it is possible for a protein to have only one homolog in human, families of singletons are not excluded.

3.1/ The query is then run on four HHpred proteomes, called “test” proteomes, corresponding to the following four species: *Arabidopsis thaliana, Drosophila melanogaster, Escherichia coli* and *Haloferax volcanii* (an archaea).

3.2/ For each species, the following step is performed:

First, the hits whose probability is greater than or equal to the previously selected threshold (see 2.1) are selected. Then, depending on their size and location on the query sequence, they are assigned, if possible, to a previously built family (see 2.2).

At the end of step 3, a family is thus made up of hits belonging at least to *Homo sapiens* and possibly to *Arabidopsis thaliana, Drosophila melanogaster, Escherichia coli* or *Haloferax volcanii*. If a family includes only human proteins, the robustness assumption can neither be rejected nor established. In this case, the threshold is lowered and step 2 is performed again.

4./ For each family, InterPro annotations (Blum *et al*., 2020) of proteins associated with hits are collected and inspected manually (in particular the part of these proteins that corresponds to the hits). If the annotations of the human proteins are similar to each other and to all proteins from at least one other organism, this family/similarity is considered “robust”; these annotations are then associated with the corresponding part of the viral protein (the query); if not, no similarities can be identified, and we consider that no similarity between the query and the human proteins can be detected.

It should be noted that when the threshold of 0.8 is reached and it is not possible to reject or establish the robustness hypothesis, an in-depth examination of the results is carried out by relaxing the constraints *i*/ on the probability threshold, which is then set to 0.5 (in accordance with the HHpred documentation which states that “typically, a match should be seriously considered if it has a probability value >50%”)), and *ii*/ on the size and location of hits; the annotations of the proteins found by relaxing the constraints are then examined; if at least 90% of human proteins are similarly annotated and these are also similarly annotated to 100% of the proteins of at least one other organism, this family/similarity is considered “robust”; these annotations are then associated with the corresponding part of the viral protein (the query).

The similarities identified at the 0.95 and 0.9 probability levels will be labeled by “highly robust”; the similarities identified at the 0.85 and 0.8 probability levels will be labeled by “very robust”; finally, the similarities identified during the relaxation stage of constraints will be labeled by “quite robust”.

Note: only proteins beginning with the prefix NP are considered in the analysis. XP records (proteins) are not curated and are therefore not considered here; furthermore, proteins identified by HHpred that do not have a match in “UniProtKB reviewed (Swiss-Prot)” (name and size in amino acids) were not considered either.

## Results

In our study, we identified a list of 6 robust similarities. We focus here on the two similarities not yet documented in the literature. For reasons of clarity, for each family/similarity considered here, only the best hit in each organism is provided. All results can be found in Supplemental file 2 (this file contains a condensed version of the results produced by HHpred which are enriched by the InterPro annotations). The raw HHpred results are stored in a separate gzip file called Supplemental file 4.

Note that the Pfam annotations of the proteins come from the InterPro or Pfam legacy (http://pfam-legacy.xfam.org/) websites; the two sites generally give similar predictions; however, the domain boundaries may sometimes differ very slightly.

### Spike S harbors a part of a “Prominin domain” (highly robust similarity)

The length of this protein is 1273 A.A.

At the 0.90 probability level, 2 human proteins share similarity with Spike S (prominin-1 and prominin-2 proteins). The best match is human prominin-1 (PROM1_HUMAN/NP_006008, length = 865). Its 186-482 part is similar to the 908-1254 part of Spike S; the 186-482 part of this human protein is included in the Pfam “Prominin” domain, Prominin/19-820. Note that when the “MSA generation iterations” parameter is set to 3 (default setting), similar results are obtained.

For the given threshold of 0.90, one fly protein annotated with the Prominin domain of Pfam shares similarities with Spike S: the fly protein “Prominin-like protein” (PROML_DROME/NP_001261351.1, length = 1013) whose 235-534 part is similar to the 911-1254 part of Spike S; the 235-534 part of this fly protein is included in the “Prominin” domain of Pfam, Prominin/76-881.

This strongly suggests that Spike S hosts part of the “Prominin domain”.

### ORF3a has similarities with some “G Protein-Coupled Receptors” (quite robust similarity)

The length of this protein is 275 A.A.

At the 0.80 probability level, a human protein shares similarity with ORF3a, the human “lutein-choriogonadotropic hormone receptor” (LSHR_HUMAN/NP_000224, length = 699). Its 537-693 part is similar to the 41-183 part of ORF3a. A large part of this 537-693 region is included in the “7 transmembrane receptor (rhodopsin family)” Pfam domain, *i*.*e*. 7tm_1/376-623. Note that when the “MSA generation iterations” parameter is set to 3 (default setting), no significant results are obtained (the probability of the best hit is 0.66).

For the given threshold of 0.80, no similarity is detected with proteins belonging to the 4 “test” proteomes. However, a number of factors support this similarity when certain constraints are relaxed (see Materials and methods):

Looking at the list of hits found by HHpred between ORF3a and the human proteome (see Supplemental file 2), it is immediately obvious that the vast majority of human proteins found are G Protein-Coupled Receptors (GPCRs). Indeed, it appears that out of 28 hits, 26 concern GPCRs (26/28 = 0.928), while the other two correspond to transmembrane segments of proteins that are not linked to GPCRs.

Considering the fly proteome and applying the same methodology as previously used in human, it appears that out of 3 hits, 3 concern GPCRs (see Supplemental file 2).

Overall (see Materials and Methods), this suggests that the similarity found is quite robust and that ORF3a shares similarities with human GPCRs.

## Discussion

The documented loss of sensitivity of Pfam HMMs when searching for domains in “divergent organisms” (Terrapon *et al*., 2012) prompted us to use HHpred (Gabler *et al*., 2020) to annotate SARS-CoV-2 proteins. Given a query sequence, this annotation tool offers the possibility to search for homologs among all proteins in an organism. Each protein in the organism is represented by an HMM built according to a different strategy than that used by Pfam (for more details, see the section “Creating custom databases” in the user guide (https://github.com/soedinglab/hh-suite/wiki)). We speculated that this difference might give HHpred the ability to discover similarities not detectable by Pfam (it should be noted that a theoretical comparison between the Pfam and HHpred HMMs, as well as a full empirical comparison, is beyond the scope of this paper).

To avoid as much as possible false predictions when using HHpred, we decided to disable its first step which is based on an iterative search strategy. Indeed, the greater the number of search iterations, the greater the risk of recruiting non-homologous sequences in the following iterations (see Materials and Methods). Furthermore, in addition to the probability assigned by HHpred to each hit, we decided to evaluate the robustness of these latter. Our evaluation procedure is based on two unimplemented ideas described in (Gabler *et al*., 2020) and can be summarized as follows (see Materials and Methods for more details; see also Figure):

A probability threshold is set; the starting value is 0.95 (according to (Gabler *et al*., 2020), when the probability of a hit is greater than 95%, homology is highly probable). Each viral protein (“query” sequence) is compared to the human proteome using HHpred; all hits with a probability above the chosen threshold are selected (if no hit meets this criterion, the threshold is successively lowered to 0.9, 0.85 and 0.80); if all hits of similar size located at the same position on the query sequence (*i*.*e*., a family of homologous hits) are annotated with the same InterPro domain (Blum *et al*., 2020), their probability of actually being homologous to the query is very high (“Check relationships among top hits”, first idea from (Gabler et al., 2020)); the query is then run on a set of “test” proteomes to check whether similarly annotated homologous hits are returned (“Check if you can reproduce the results with other parameters”, second idea of (Gabler *et al*., 2020)); if so, a family of homologous hits defined a “robust similarity”; if not, we consider that no similarities can be identified. Note that when a family includes only human proteins, the robustness assumption can neither be rejected nor established; in this case, the threshold is lowered and the study is carried out again. It should be also noted that when the threshold of 0.8 is reached and it is not possible to reject or establish the robustness hypothesis, a thorough examination of the results is carried out by relaxing the constraints (mainly on the size, location and/or probability associated with the hits, see Materials and Methods for more details). Similarities identified at the 0.95 and 0.9 probability levels are labeled “highly robust”; similarities identified at the 0.85 and 0.8 probability levels are labeled “very robust”; finally, the similarities identified when certain constraints are relaxed are described as “quite robust”.

The organisms used to evaluate the HHpred results are *Arabidopsis thaliana, Drosophila melanogaster, Escherichia coli* and *Haloferax volcanii* (an archaea). Note that, in order to potentially increase the identified similarities, we would have liked to include proteomes from organisms closer to humans in our study. Unfortunately, the online server currently does not offer the option to use such proteomes. To successfully accomplish this task, it is necessary to perform the local installation of the free HH-suite software and build these proteomes using this software. This work needs to be done (future works).

Below we present a summary of our results.

We subjected the 17 proteins of the SARS-CoV-2 proteome (see Materials & Methods and Results sections) to our annotation procedure. UniProt considers polyproteins 1a (pp1a) and 1ab (pp1ab) as two separate entries; polyprotein pp1ab is proteolytically cleaved to form 15 shorter proteins; the first 10 proteins (NSP1, …, NSP10) are also cleaved from pp1a; NSP12, …, NSP16 are unique to pp1ab. We therefore subjected 30 proteins to our evaluation procedure.

### No “robust” similarities were found for the following 24 proteins

NSP1, NSP4-10, NSP12, NSP14-15, Nucleoprotein, Envelope small membrane, Membrane Protein M, ORF3B, ORF3C, ORF3D, ORF6, ORF7a, ORF7b, ORF8, ORF9b, ORF9C, ORF10.

### A “highly robust” or “very robust” similarity, already documented in literature, was detected on the following 4 proteins

NSP3 is a papain-like protease; we showed it harbors a Macro domain. NSP13 is a helicase; we provide evidence suggesting that it harbors AAA domains. NSP16 is a methyltransferase; we confirm that it harbors a “FtsJ-like methyltransferase” domain. As these similarities are well documented, the interested reader is invited to consult the InterPro annotations.

NSP2 is involved in the inhibition of the antiviral response and facilitates SARS-CoV-2 replication. We showed that part 151-195 of NSP2, *i*.*e*. part 332-376 of polyprotein 1a, contains a “signature of the beta subunit of casein kinase II”. According to PROSITE, such a domain could be involved in the binding of a metal such as zinc. Interestingly, the structure of the N-terminal part of NSP2 was recently solved (Ma *et al*., 2021). It shows that NSP2 has three zinc fingers: Zn1, Zn2 and Zn3. Two Zn2 (resp. Zn3) binding sites are located at positions 161 and 164 (resp. at positions 190 and 193). Our prediction is therefore in agreement with this structure of the N-terminal domain of SARS-CoV-2 NSP2.

### A previously unknown “highly robust” similarity was detected on Spike S protein

The Spike S protein (1273 A.A.) is composed of two subunits: the S1 subunit (14-685 residues), and the S2 subunit (686-1273 residues), which are responsible for receptor binding and membrane fusion respectively (Huang *et al*., 2020). We have shown that the 908-1254 part of the Spike S protein is similar to the 186-482 part of human prominin-1 (length = 865). This similarity encompasses the heptapeptide repeat 1 sequence, *i*.*e*. HR1 (912-984 residues), HR2 (1163-1213 residues), the TM domain (1213-1237) and part of the cytoplasmic domain (1237-1273) of the S2 subunit; however, it excludes the fusion peptide (FP) (788-806) of S2 which plays an essential role in mediating membrane fusion. HR1 and HR2, which are part of the similarity, have been shown to form a six-helix bundle that is essential for the fusion and viral entry function of the S2 subunit (Xia *et al*., 2020).

Recently, in searching for proteins involved in SARS-CoV-2 entry into host cells, (Kotani *et al*., 2022) found that the glycoprotein CD133, the other name for prominin-1, colocalises with ACE2 – the main cell entry receptor for SARS-CoV-2 – bound to the Spike S protein in Caco-2 cells. They demonstrated that the SARS-CoV-2 Spike protein exhibited increased binding capacity in cells co-expressing ACE2 and CD133, compared to cells expressing ACE2 alone. In addition, they experimentally infected HEK293T cells with a SARS-CoV-2 pseudovirus and showed that infectivity was twice as high in HEK293T cells co-expressing CD133-ACE2 than in HEK293T cells expressing ACE2 alone. They concluded that CD133, although not a primary receptor for the SARS-CoV-2 Spike protein, is a cofactor (a co-receptor) that partially contributes to infection in the expressing cells. All these results suggest that the C-terminal part of Spike S, which has similarities with prominin-1, may be involved in the docking of Spike S to ACE2 (insofar as CD133 enhances the ability of Spike S to bind to ACE2). This obviously remains to be demonstrated but is clearly an interesting avenue of research.

While considerable work has been done to characterise the cellular receptors and pathways mediating virus internalisation, little is known about the onset of the infection process, which begins when the virus comes into contact with the host cell surface; some studies have shown that viruses “diffuse” onto the surface of host cells after “landing” on them; this process ranges from a random walk to a constrained diffusion where the virus particles appear to be confined to a specific microdomain of the cell membrane (Boulant *et al*., 2015). From this point of view, it is interesting to note that it was recently shown by (Rouaud *et al*., 2022) that *i*/ ACE2 concentrates at epithelial apical cell junctions in cultured epithelial cell lines, and that *ii*/ (Pinto *et al*., 2022) showed that ACE2 and TMPRSS2 (which is used by SARS-CoV-2 for Spike S-protein priming (Hoffmann *et al*., 2020)) were localised at the plasma membrane, including the microvilli, in human airway epithelium. Interestingly, about 25 years ago, prominin was shown to be localised to the apical surface of various epithelial cells, where it is selectively associated with microvilli and microvillus-related structures (Weigmann *et al*., 1997). Furthermore, Weigmann and colleagues showed that prominin expressed ectopically in non-epithelial cells was also selectively found in microvillus-like protrusions of the plasma membrane. Two years later, (Corbeil *et al*., 1999) showed that prominin contains dual targeting information, for direct delivery to the apical domain of the plasma membrane and for enrichment in the microvilli subdomain. Furthermore, they showed that this dual targeting does not require the cytoplasmic C-terminal tail of prominin (*i*.*e*., part 814-865 of CD133). From the above results, it is tempting to assume that the prominin-like part of Spike S is involved in the delivery of the virus to the apical domain of the plasma membrane where the ACE2 proteins are located. This hypothesis is all the more tempting as the similarity between Spike S and prominin does not concern the C-terminal part of prominin, which, as we have pointed out above, is not necessary for prominin targeting (recall that we have shown that the 186-482 part of human prominin-1 is similar to the 908-1254 part of Spike S). Unfortunately, to date, the molecular nature of the prominin apical sorting signal is unknown. It has been suggested in (Weigmann *et al*., 1997) that prominin may interact with the actin cytoskeleton, or that plasma membrane protrusions may have a specific lipid composition/organisation for which prominins may have a preference.

Finally, it should be noted that the “SARS-CoV(−1)” glycoprotein Spike, which, like SARS-CoV-2 Spike, binds to human ACE2 (Li *et al*., 2003), is also similar to human prominin-1. Specifically, using HHpred, we showed that the 177-473 part of the latter is similar to the 890-1236 part of Spike (with an associated probability of 0.95 – see Supplemental file 4, raw HHpred data). In contrast, the MERS-CoV Spike glycoprotein (like SARS-CoV and SARS-CoV-2, MERS-CoV is a betacoronavirus), which uses human DPP4 as an entry receptor (Raj *et al*., 2013), is similar to human mucin-1: the 292-421 part of mucin-1 is similar – with an associated probability of 0.89 – to the 1230-1344 part of MERS-CoV Spike (see Supplemental file 4, raw HHpred data). It is also interesting to note that (Kotani *et al*., 2022) showed that the DPP4 protein also colocalises with ACE2 and CD133 in Caco-2 cells. This suggests that it is likely that *i*/ different coronaviruses compete at the same positions on the cell, but *ii*/ use different entry receptors and therefore different types of spike proteins to reach these sites and fuse with the cells.

### A previously unknown “quite robust” similarity was detected on ORF3a protein

The 41-183 part of ORF3a (275 A.A.) shows similarities to human G Protein-Coupled Receptors (GPCRs) (which are cell surface receptor proteins that detect molecules from outside the cell and trigger cellular responses (Lagerström & Schiöth, 2008)) and in particular to the GPCRs annotated with the Pfam domain “7 transmembrane receptor (rhodopsin family)/7tm_1” (see Results section and Supplemental file 2). According to Pfam, this family contains, among other GPCRs, members of the opsin family, which are considered typical members of the rhodopsin superfamily.

The ORF3a protein of “SARS-CoV(−1)” has been shown to form an ion channel (Lu *et al*., 2006). Recently, (Kern *et al*., 2021) presented Cryo-EM determined structures of SARS-CoV-2 ORF3a at a resolution of 2.1Å. The authors provide evidence suggesting that ORF3a forms a large polar cavity in the inner half of the transmembrane region (TM) that could form ionic conduction paths (TM1 (43-61), TM2 (68-99) and TM3 (103-133)). Interestingly, the similarity we detected on ORF3a (41-183) encompasses the transmembrane portion of ORF3a (43-133) which could form ionic permeation pathways. As mentioned earlier, we have shown that this part of ORF3a resembles many GPCRs which belong to the Rhodopsin family (22 of 28 human proteins sharing similarities with ORF3a, see Supplemental file 2 for more details). It is interesting to note that some GPCRs, called “Rhodopsin channels”, directly form ion channels (see (Nagel *et al*., 2002) and (Nagel *et al*., 2003)). From this point of view, our prediction is therefore in line with the work of (Kern *et al*., 2021). However, it is worth mentioning that a recent work challenges the results of both (Kern *et al*., 2021) and (Lu *et al*., 2006): (Miller *et al*., 2023) provide evidence suggesting that while a narrow cavity is detected in the SARS-CoV-2 ORF3a transmembrane region, it likely does not represent a functional ion-conducting pore (the same holds true for SARS-CoV-1 ORF3a).

Finally, it should be noted that if our method is applied to the ORF3a of SARS-CoV(−1), no similarities are identified. More precisely, none of the similarities found by HHpred are significant, *i*.*e*. the probability of the best hit is 0.72, which is below our threshold of 0.8; moreover, this best hit does not correspond to a GPCR (see Supplemental file 4). This result may suggest a lack of sensitivity of HHpred. That said, although HHpred is a fairly effective tool for detecting very distant homologies, not all similarities are detectable. Furthermore, although the ORF3a of SARS-CoV(−1) and SARS-CoV-2 share 72% sequence identity and are similar in the arrangement of the TM domains, the differences observed in the ion channel properties between these two proteins suggest a different mode of action between them (Zhang *et al*., 2022).

### Comparison of our results with those of “Pfam clans”

As indicated in the introduction to this article (see also Supplemental file 1), of the 40 Pfam domains that annotate SARS-CoV-2 proteins, only one domain is not confined to viruses, the Macro domain that annotates NSP3. This observation can be modulated at the level of Pfam clans which are collections of related domains. At this level, 12 domains belong to clans whose domains are not strictly viral (see Supplemental file 1). These clans allow the annotation of the following 9 proteins (more generally, of only part of each protein): NSP3, NSP5, NSP13, NSP14, NSP15, NSP16, ORF7a, ORF8, and Spike S. 4 of these proteins are annotated by both Pfam and our approach: NSP3, NSP13, NSP16 and Spike S. In the case of NSP3, NSP13 and NSP16, the annotations are similar (note however that for NSP3, Pfam detects two domains related to the MACRO clan; only one Macro domain is detected by our approach) whereas in the case of Spike S, our annotations refer to a different part of the protein than that annotated by Pfam. We also identified similarities, not restricted to viruses unlike Pfam, for ORF3a and NSP2.

### Evaluation of our results in light of the known weaknesses of HHpred

As reported in (Gabler *et al*., 2020) and (Kuchibhatla *et al*., 2014), some false positive HHpred hits may have high scores because they have coiled-coil, transmembrane or low complexity segments. Of our 6 “robust similarities”, 2 have transmembrane segments and/or disordered areas (according to InterPro annotations).

#### ORF3a

As previously indicated, ORF3a shares similarity with G Protein-Coupled Receptors (GPCRs) annotated with the Pfam domains “7 transmembrane receptor (rhodopsin family)/7tm_1” or “7 transmembrane receptor (secretin-like) 7tm_2” (see Results or Supplemental file 2).

Since transmembrane proteins are a large family of proteins – according to UniProt, out of 80581 proteins expressed by humans, 13876 are transmembrane proteins – it is legitimate to ask whether the (observed) distribution of transmembrane proteins found by HHpred – out of 28 proteins found by HHpred, 28 are transmembrane proteins – is the same as the (expected) distribution of transmembrane proteins in UniProt. Using a Fisher’s exact test, we conclude (see Supplemental file 3 for proof) that the results found by HHpred are not randomly drawn from the UniProt human proteome (p-value = 6.2059249716913E-11).

Similarly, as transmembrane proteins can be grouped into many different classes (the Pfam clan “Family A G protein-coupled receptor-like superfamily”, to which 7tm_1 and 7tm_2 belong, alone contains 53 different domains), it can also be argued that the similarities found by HHpred are due to chance. Of the 28 transmembrane proteins found by HHpred, 26 belong to the 7tm_1 or 7tm_2 classes. Knowing that the number of human proteins belonging to the 7tm_1 or 7tm_2 classes is – according to UniProt – 540, we show (see Supplemental file 3 for proof) using a Fisher’s exact test that the results obtained by HHpred do not arise from random selection within the different classes of the transmembrane protein family (p-value = 2.8739559680731E-12).

#### Spike glycoprotein

As shown previously, the 908-1254 part of the Spike S protein of SARS-CoV-2 is similar to the 186-482 part of human prominin-1. The 179-432 part of this prominin is annotated as “NON_CYTOPLASMIC_DOMAIN” (*i*.*e*. non-cytoplasmic loops of a TM protein) by Phobius (for completeness, note that the 253-283 part is annotated as a coil by COILS).

In contrast to the case of ORF3a, no reliable statistical test can be performed here (the number of human prominins, *i*.*e*. proteins annotated by Pfam as “prominin” (Pfam PF05478), is 5). However, such a calculation seems unnecessary here. HHpred identified a similarity between Spike S and human and fly prominins (see Results section). Human and fly belonging to lineages that were separated over 700 million years ago (median time of divergence 694 MYA (see http://timetree.org/, (Kumar et al., 2017)), this similarity is clearly not a coincidence (unless one imagines a recent horizontal transfer).

### Further investigation of already found similarities

In the previous sections, we provided evidence suggesting that SARS-CoV-2 Spike S shares similarities with Human Prominin, with Prominin 1 as the closest match, and ORF3a with Human G-coupled proteins, with “Lutropin-choriogonadotropic hormone receptor” as the closest match. To further investigate these similarities, we decided to check whether Prominin 1 (resp. Lutropin-choriogonadotropic hormone receptor) resembles other viral proteins and, if so, which ones. Using HHPRED (default parameters), we ran Prominin 1 (resp. Lutropin-choriogonadotropic hormone receptor) against a database containing all viral protein sequences from SwissProt as of November 2021 (keep in mind that SARS-CoV-2 proteins are not included in this dataset).

Very interestingly, the analysis of the results obtained by HHPRED on Human Prominin 1 revealed that the part of Human Prominin 1 that shares similarities with SARS-CoV-2 Spike S (*i*.*e*., part 186-482 of Prominin 1 is similar to the part 908-1254 of Spike S) also shares similarities with 6 viral Spike glycoproteins (with a probability greater than 0.95), all of which are expressed by coronaviruses (Human coronavirus NL63, Human coronavirus 229E, Bat coronavirus 512/2005, Porcine epidemic diarrhea virus (which is a coronavirus), Canine coronavirus (strain Insavc-1), Murine coronavirus (strain A59)) (see Supplemental file 5).

The results obtained by HHPRED on the “Lutropin-choriogonadotropic hormone receptor” are equally interesting but more challenging to interpret directly. On the one hand, they clearly show that the part of this G-coupled protein that shares similarities with SARS-CoV-2 ORF3a (*i*.*e*., part 537-693 of “Lutropin” is similar to part 41-183 of ORF3a) shares similarities with 23 viral proteins - with a probability greater than 0.99 -, all of which are annotated as “G-protein coupled receptors” (see Supplemental file 5). However, on the other hand, the detected similarities encompass the region of interest in “Lutropin” (*i*.*e*., part 537-693), which, as a subset of the overall similarities, cannot be directly associated with the computed probability. To address this issue, we re-ran HHPRED using only the 537-693 part of the Lutropin-choriogonadotropic hormone receptor. HHPRED returns the same 23 proteins associated with a probability match greater than 0.98 (see Supplemental file 5). Note that G-protein coupled receptors belonging to the “Herpesvirales” order are the most represented (17 occurrences), including various strains of Human herpesvirus, Equine herpesvirus, Elephantid herpesvirus, Murid herpesvirus, Saimiriine herpesvirus, and Alcelaphine herpesvirus, as well as Epstein-Barr virus and Human cytomegalovirus which are also herpesviruses.

These results thus support the predictions made using our methodology. The overall process can be viewed as a reciprocal best hit (or bidirectional best hit): 1. a part of SARS-CoV-2 Spike S is similar to a part of human Prominin 1, which is itself similar to several viral Spike glycoproteins; 2. a part of SARS-CoV-2 ORF3a is similar to a part of a human G-coupled protein, which is also itself similar to several viral G-coupled proteins.

## Conclusion

We used HHpred to search for similarities between SARS-Cov-2 and human proteins. To avoid false predictions, the robustness of each similarity was assessed using a procedure based on “test sets/proteomes”. We found six robust similarities in six different proteins, of which three are already documented, one is in agreement with recent crystallographic results, and two are not reported in the literature. We focused on these last two similarities and showed how they open new avenues of research to better understand this virus. Obviously, our work is limited to making predictions that need to be validated experimentally. Furthermore, the origin of the similarities (evolutionary convergence, horizontal transfer, etc.) has not been addressed in this work. Nevertheless, we believe that our approach (or one similar to it) can be profitably used to open up lines of research and to improve the annotation of any virus, especially “orphan viruses”, *i*.*e*. viruses which, for various reasons, are far much less studied than SARS-CoV-2.

## Acknowledgements

We would like to thank all those who initiated Pfam, InterPro and HH-suit and all those who maintain and improve these databases and tools for the benefit of our community. Without them, the work presented here would not have been possible. We also warmly thank Joël Pothier (ISYEB, MNHN), the PCI Genomics editor of this paper, and the two anonymous referees whose comments have clearly contributed to improve the quality of this paper.

## Supplementary information availability

Supplemental files 1, 2, 3 and 4 are available online (*cf*. bioRxiv, DOI of this document).

