## Supplemental file 1 for "Re-annotation of SARS-CoV-2 proteins using an HHpred-based approach opens new opportunities for a better understanding of this virus"

### Pfam annotation of SARS-CoV-2 proteins

The SARS-CoV-2 virus expresses 17 proteins, see UniProt (https://www.uniprot.org/); UniProt provides polyproteins 1a (pp1a) and 1ab (pp1ab) as two separate entries. The pp1ab polyprotein is cleaved to form 15 shorter proteins; the first 10 proteins, *i.e.*, NSPs 1-10, are also cleaved from pp1a; NSPs 12-16 are unique to pp1ab. When we look at the Pfam annotations of these proteins (see below), we see that 4 of these 17 proteins are not Pfam annotated (these 4 proteins are small, *i.e.* between 22 and 57 amino acids); the other 13 proteins are annotated with a set of 40 domains, 39 of which are strictly associated with viruses (the Macro domain is the exception to the rule) This shows very clearly that the SARS-CoV-2 proteins are essentially virus-like (97.5% (39/40) of the annotated domains of this virus are virus-restricted).

The above observation can be modulated at the level of Pfam clans which are collections of related domains. At this level, 12 domains belong to clans whose domains are not strictly viral. Under these conditions, 70% (1-12/40) of the SARS-CoV-2 domains resemble viral domains about which little information is available.

Note : the Pfam annotations of the proteins come from the InterPro or Pfam legacy (http://pfam-legacy.xfam.org/) websites; the two sites generally give similar predictions; however, the domain boundaries may sometimes differ very slightly.

**« Accessory proteins » (*i.e.*, all proteins expressed by SARS-CoV-2 except polyproteins 1a and 1ab) :**

Summary: 15 proteins; 4 unannotated Pfams; 14 virus-restricted Pfam domains; 4 domains are members of a Pfam clan.

A0A663DJA2_SARS2:
Length:38
No Pfam annotation.

AP3A_SARS2 :
Length:275
Pfam bCoV_viroporin/ 1-274
Widespread in viruses (Orthornavirae).

NCAP_SARS2 :
Length:419
Pfam CoV_nucleocap/21-391
Widespread in viruses (Orthornavirae)

NS8_SARS2 :
Length:121
Pfam bCoV_NS8/1-118
Widespread in viruses (Orthornavirae).
This family is a member of clan Ig (CL0011)

NS7B_SARS2 :
Length:43
Pfam bCoV_NS7B/1-43
Widespread in viruses (Orthornavirae)

NS6_SARS2 :
Length:61
Pfam bCoV_NS6/1-61
Widespread in viruses (Orthornavirae)

SPIKE_SARS2 :
Length:1,273
Pfam bCoV_S1_N/29-337
Pfam bCoV_S1_RBD/348 -526
Pfam CoV_S1_C/536-594
Pfam CoV_S2/710-1232

bCoV_S1_N :
Widespread in viruses (Orthornavirae)
This family is a member of clan Concanavalin (CL0004)

bCoV_S1_RBD :
Widespread in viruses (Orthornavirae)

CoV_S1_C :
Widespread in viruses (Orthornavirae)

CoV_S2:
Widespread in viruses (Orthornavirae)
This family is a member of clan Fusion_gly (CL0595)

NS7A_SARS2 :
Length:121
Pfam bCoV_NS7A/16-121
Widespread in viruses (Orthornavirae)
This family is a member of clan Ig (CL0011)

ORF3B_SARS2 :
Length:22
No Pfam annotation

ORF3C_SARS2 :
Length:41
No Pfam annotation

ORF3D_SARS2 :
Length:57
No Pfam annotation

ORF9B_SARS2 :
Length:97
Pfam bCoV_lipid_BD /1 -97
Widespread in viruses (Orthornavirae)

ORF9C_SARS2 :
Length:73

Pfam  bCoV_Orf14/1-70
Widespread in viruses (Orthornavirae)

VEMP_SARS :
Length:   76
Pfam  CoV_E /2-76
Widespread in viruses (Orthornavirae)

VME1_SARS2 :
Length:222
Pfam  CoV_M/15-218
Widespread in viruses (Orthornavirae)

**Polyprotein 1a and 1ab (1-4405 + 4406-7096) :**
Lenght : 7096

Summary : 26 Pfam domains ; 25 restricted to virus and one widespread in eukaryotes, bacteria and viruses (Macro) ; 9 domains are members of a Pfam clan ; one of these 9 clans is restricted to viruses (CL0027).

NSP1 :

Pfam  bCoV_NSP1/ 7-144
Widespread in viruses (Orthornavirae +)

NSP2 :

Pfam  CoV_NSP2_N/182-423
Widespread in viruses (Orthornavirae +)

Pfam  CoV_NSP2_C/652-818
Widespread in viruses (Orthornavirae +)

NSP3 :

Pfam  bCoV_NSP3_N/880-1051
Widespread in viruses (Orthornavirae +)

Pfam  Macro/1058-1165
Widespread in eukaryotes, bacteria and viruses
This family is a member of clan MACRO  (CL0223)

Pfam  bCoV_SUD_M/1351-1493
Widespread in viruses (Orthornavirae +)
This family is a member of clan MACRO (CL0223)

Pfam  bCoV_SUD_C/1497-1561
Widespread in viruses (Orthornavirae +)

Pfam  CoV_peptidase/1564-1882
Widespread in viruses (Orthornavirae +)
This family is a member of clan Peptidase_CA (CL0125)

Pfam bCoV_NAR/1909-2019
Widespread in viruses (Orthornavirae +)

Pfam CoV_NSP3_C/2260-2749
Widespread in viruses (Orthornavirae +)

NSP4 :

Pfam CoV_NSP4_N/2788-3142
Widespread in viruses (Orthornavirae +)

Pfam CoV_NSP4_C/3166-3262
Widespread in viruses (Orthornavirae +)

NSP5 :

Pfam Peptidase_C30/3292-3582
Widespread in viruses (Orthornavirae +)
This family is a member of clan Peptidase_PA (CL0124)

NSP6 :

Pfam CoV_NSP6/3597-3859
Widespread in viruses (Orthornavirae +)

NSP7 :

Pfam CoV_NSP7 /3860-3942
Widespread in viruses (Orthornavirae +)

NSP8 :

Pfam CoV_NSP8/3943-4139
Widespread in viruses (Orthornavirae +)

NSP9 :

Pfam CoV_NSP9/4141-4253
Widespread in viruses (Orthornavirae +)

NSP10 :

Pfam CoV_NSP10/4262-4384
Widespread in viruses (Orthornavirae +)

///

NSP12 :

Pfam CoV_Rpol_N/4406-4758
Widespread in viruses (Orthornavirae)

Pfam RdRP_1/4858-5195
Widespread in viruses
This family is a member of clan RdRP (CL0027) (viruses)

NSP13 :

Pfam Viral_helicase1/5634 -5900
Widespread in viruses, bacteria and arthropoda
This family is a member of clan P-loop_NTPase (CL0023)

NSP14 :

Pfam CoV_ExoN/5928-6450
Widespread in viruses (Orthornavirae +)
This family is a member of clan NADP_Rossmann (CL0063)

NSP15 :

Pfam CoV_NSP15_N/6453-6513
Widespread in viruses (Orthornavirae +)

Pfam CoV_NSP15_M/6514-6637
Widespread in viruses (Orthornavirae +)

Pfam CoV_NSP15_C/6643-6796
Widespread in viruses (Orthornavirae +)
This family is a member of clan EndoU (CL0695)

NSP16 :

Pfam CoV_Methyltr_2/6800-7096
Widespread in viruses (Orthornavirae +)
This family is a member of clan NADP_Rossmann (CL0063)

Considering that the SARS-CoV-2 domains that belong to a clan (i.e. a superfamily of related domains) comprising domains that are widespread in eukaryotes, 70% (28 out of 40) of the SARS-CoV-2 domains are still without homologs outside of the viral taxa.
