## Supplemental file 2 for "Re-annotation of SARS-CoV-2 proteins using an HHpred-based approach opens new opportunities for a better understanding of this virus"

### Results

In the following sections, we present key findings of our analysis. These results are a summary of the outputs produced by HHpred (see Supplemental file 4).

Note that the Pfam annotations of the proteins come from the InterPro or Pfam legacy (http://pfam-legacy.xfam.org/) websites; the two sites generally give similar predictions; however, the domain boundaries may sometimes differ very slightly.

<u>NSP2 harbors a “Casein kinase II regulatory subunit pattern” (very robust similarity)</u>

NSP2 derives from polyprotein 1a (181-818). The length of this protein is 638 A.A.

At the 0.85 probability threshold, only one human protein shares similarity with NSP2. Specifically, the 151-195 part of NSP2 is similar to the 101-142 part of the human protein "Casein kinase II subunit beta" (CSK2B_HUMAN/NP_001311, length = 215 A.A.). The 109-140 part of the human "Casein kinase II subunit beta" protein is annotated with the PROSITE "Casein kinase II regulatory subunit signature" motif ; in addition, parts 105-126 and 127-148 of this protein are annotated with the PRINTS motif "CASNKINASEII". This suggests that the 151-195 part of NSP2 could also be a "casein kinase II regulatory subunit signature". Note that when the "MSA generation iterations" parameter is set to 3 (default setting), no significant results are obtained (the probability of the best hit is 0.32).

For the given threshold of 0.85, this part of NSP2 is also similar to a part of an *Arabidopsis thaliana* protein. Specifically, the 151-194 part of NSP2 is similar to the 182-222 part of the *Arabidopsis thaliana* protein "Casein kinase II subunit beta" (CSK2D_ARATH/NP_191584.1, length = 276 AA). The 190-221 part of the latter is annotated with the PROSITE motif "Casein kinase II regulatory subunit signature" ; in addition, parts 186-207 and 208-229 of this protein are annotated with the PRINTS motif "CASNKINASEII".

This  suggest that NSP2 harbours a “Casein kinase II regulatory subunit pattern”.

<u>NSP3 harbors a Macro domain (highly robust similarity)</u>

NSP3 derives from polyprotein 1a (819-2763). The lenght of this protein is 1945 A.A.

For the 0.95 probability threshold, 7 human proteins share similarity with NSP3. The best result is the human "Core histone macro-H2A.2" protein (H2AW_HUMAN/NP_061119, length = 372) whose 187-371 part is similar to the 210-377 part of NSP3, *i.e.* the 1029-1197 part of polyprotein 1a. The 187-371 region of this human protein contains the Pfam Macro domain (216-329). This suggests that the 210-377 part of NSP3 also shares similarity with Macro. Note that when the "MSA generation iterations" parameter is set to 3 (default setting), similar results are obtained (so, for example, the best hit with the default setting is the fourth best hit when "MSA generation iterations" parameter is set to 0).

The following six human matches are consistent with the above findings:

Protein mono-ADP-ribosyltransferase PARP14 (PAR14_HUMAN/NP_060024, length = 1801) : 219-375  NSP3 <=> 803-977, Macro/820-936.

ADP-ribose glycohydrolase MACROD2 (MACD2_HUMAN/NP_542407, length = 425) : 215-373 NSP3 <=> 67-240, Macro/88-200.

Protein mono-ADP-ribosyltransferase PARP9 (PARP9_HUMAN/NP_001139574, length = 854) : 219-375 NSP3 <=> 119-295, Macro/136-253.

ADP-ribose glycohydrolase MACROD1 (MACD1_HUMAN/NP_054786, length = 325) : 209-372 NSP3 <=> 143-318, Macro/170-282.

Protein mono-ADP-ribosyltransferase PARP15 (PAR15_HUMAN/NP_001106995, length = 678) : 219-377 NSP3 <=> 90-268, Macro/107-220.

Chromodomain-helicase-DNA-binding protein 1-like (CHD1L_HUMAN/NP_004275, length = 897) : 217-362 NSP3 <=> 714-877, “Macro domain”/704-897 (according to InterPro).

For the probability threshold considered (i.e., 0.95), one *Escherichia coli* protein shares similarity with NSP3: the "O-acetyl-ADP-ribose deacetylase" protein (YMDB_ECOLI/NP_415563, length = 177) whose 3-166 part is similar to the 218-367 part of NSP3; the 218-367 region of this bacterial protein contains the Pfam Macro domain (21-137).

Previous elements strongly suggests that NSP3 harbors a Macro domain.

<u>NSP13 harbors AAA domains (highly robust similarity)</u>

NSP13 derives from Polyprotein 1ab (5325-5925). The length of this protein is 601 A.A.

At the 0.95 probability level, many human proteins share similarity with NSP13. The best match is the human SMUBP-2 protein (SMBP2_HUMAN/NP_002171, length = 993) whose 207-618 part is similar to the 275-582 part of NSP13. The 207-618 part of this human protein is involved in two Pfam domains, namely AAA_11/191-411 and AAA_12/418-615, which are both members of the P-loop NTPase clan (CL0023). This suggests that the 275-582 part of NSP13, i.e. the 5600-5907 part of polyprotein 1ab, harbours AAA domains. Note that when the "MSA generation iterations" parameter is set to 3 (default setting), similar results are obtained (in this case, for example, the first best hit is the same).

The following 14 best matches are consistent with the previous results:

Helicase MOV-10 (MOV10_HUMAN/NP_001123551, length = 1003) : 280-581 NSP13 <=> 522-925, AAA_30/ 498-696 + AAA_12/698-923.

DNA replication ATP-dependent helicase/nuclease DNA2 (DNA2_HUMAN/NP_001073918, lenght = 1060) : 275-581 NSP13 <=> 641-1020, AAA_11/625-736 + AAA_11/722-798 + AAA_12/805-1017.

Regulator of nonsense transcripts 1 (RENT1_HUMAN/NP_001284478.1, lenght = 1129): 275-581 NSP13 <=> 496-890, InterPro P-loop_NTPase (P-loop containing nucleoside triphosphate hydrolase)/445-925 ; AAA_11/584-682 + AAA_12/691-887.

RNA helicase Mov10l1 (M10L1_HUMAN/NP_061868.1, lenght = 1211): NSP13 280-581 <=> 758-1152, InterPro P-loop_NTPase (P-loop containing nucleoside triphosphate hydrolase)/663-1176 ; AAA_11/745-827 + AAA_11/856-926 + AAA_12/937-1149.

Probable helicase senataxin (SETX_HUMAN/NP_055861, lenght = 2677) : 276-581 NSP13 <=> 1957-2428, AAA_11/856-926 + AAA_12/2225-2426.

Helicase with zinc finger domain 2 (HELZ2_HUMAN/NP_001032412.2, length = 2649) : 275-581 NSP13 <=> 2167-2614, AAA_11/2152-2398 + AAA_12/2406-2611.

Protein ZGRF1 (ZGRF1_HUMAN/NP_060862, lenght = 2104) : 276-581 NSP13 <=> 1648-2043, AAA_11/1757-1847 + AAA_12/1855-2041.

F-box DNA helicase 1 (FBH1_HUMAN/NP_835363.1, lenght = 1043) : 280-581 NSP13 <=> 461-938, InterPro P-loop_NTPase (P-loop containing nucleoside triphosphate hydrolase)/433-946 ; AAA_19/457-681 + UvrD_C (member of the P-loop_NTPase clan)/869-934.

RNA helicase aquarius (AQR_HUMAN/NP_055506, length = 1485) : 280-581 NSP13 <=> 821-1310, AAA_11/801-1107 + AAA_12/1115-1308.

Schlafen family member 5 (SLFN5_HUMAN/NP_659412.3, length = 891) : 280-577 NSP13 <=> 576-898, P-loop_NTPase/539-891.

NFX1-type zinc finger-containing protein 1 (ZNFX1_HUMAN/NP_066363.1, length = 1918) : NSP13 275-581 <=> 612-1237, P-loop_NTPase/556-1261.

DNA helicase B (HELB_HUMAN/NP_387467, length = 1087) : NSP13 275-581 <=> 469-937, P-loop_NTPase/303-948.

Protein SLFN14 (SLN14_HUMAN/NP_001123292, length = 912) : NSP13 276-582 <=> 587-912, P-loop_NTPase/576-709.

ATP-dependent DNA helicase PIF1 (PIF1_HUMAN/NP_079325.2, length = 641) : NSP13 280-581 <=> 226-600, P-loop_NTPase/188-614.

The previously considered part of NSP13 is similar to three (reviewed) *Arabidopsis thaliana* proteins. The best match is the Arabidopsis thaliana "probable helicase” protein (MAA3_ARATH/NP_001329005, length = 818) whose 273-734 part is similar to the 275-581 part of NSP13. The 273-734 part of this plant protein is involved in two Pfam domains, namely AAA_11/257-436 + AAA_11/451-526 + AAA_12 /534-731, which are members of the P-loop NTPase clan (CL0023). This result is in agreement with what has been found in human.

The following two matches are also consistent with the above:

Regulator of nonsense transcripts 1 homolog (RENT1_ARATH/NP_199512) : 275-581 NSP13 <=> 503-899, InterPro P-loop_NTPase (P-loop containing nucleoside triphosphate hydrolase)/452-934 ; AAA_11/591-691 + AAA_12/701-896.

DNA replication ATP-dependent (JHS1_ARATH/NP_001322072.1) : 275-581 NSP13 <=> 938-1302, AAA_11/923-1017 + AAA_11/1023-1090 + AAA_12/1100-1299.

Our results strongly suggest that the part 275-582 of NSP13 harbors AAA domains.

NSP16 is a methyltransferase (highly robust similarity)

NSP16 derives from Polyprotein 1ab (6799-7096). The length of this protein is 298 A.A.

At the 0.95 probability level, two human proteins share similarity with NSP16. The best match is the "pre-rRNA 2'-O-ribose RNA methyltransferase FTSJ3" (SPB1_HUMAN/NP_060117, length = 847). Its 31-217 part is similar to the 46-230 part of NSP16, i.e. the 6845-7029 part of polyprotein 1ab. The 31-217 part of this human protein corresponds quite well to the Pfam "FtsJ-like methyltransferase" domain, FtsJ/21-207. The other match is the "Putative tRNA (cytidine(32)/guanosine(34)-2'-O)-methyltransferase" (TRM7_HUMAN/NP_036412.1, length = 329) whose 28-203 part is similar to the 46-215 part of NSP3. The 28-203 part of this human protein again corresponds quite well to the Pfam "FtsJ-like methyltransferase" domain, FtsJ/21-207. Note that when the "MSA generation iterations" parameter is set to 3 (default setting), similar results are obtained (thus, for example, the best result is the same in both cases).

The part of NSP16 considered above is similar to two proteins from Drosophila melanogaster. The best match is the fly protein "Putative tRNA (cytidine(32)/guanosine(34)-2'-O)-methyltransferase 1" (TRM71_DROME/NP_650590, length = 302) whose 28-211 part is similar to the 46-215 part of NSP16. The 28-211 part of this human protein corresponds quite well to the Pfam "FtsJ-like methyltransferase" domain, FtsJ/21-207. The second match is the "Putative tRNA (cytidine(32)/guanosine(34)-2'-O)-methyltransferase 2" protein (TRM72_DROME/NP_650947.1, length = 320) whose 28-209 part is similar to the 46-215 part of NPS16. The 28-209 part of this human protein again corresponds quite well to the Pfam FtsJ domain, i.e. FtsJ/21-205. This result is in agreement with what has been found in humans.

This suggests that NSP16 is a methyltransferase.

Spike glycoprotein harbors a part of a “Prominin domain” (highly robust similarity)

At the 0.90 probability level, 2 human proteins share similarity with Spike S (prominin-1 and prominin-2 proteins). The best match is human prominin-1 (PROM1_HUMAN/NP_006008, length = 865). Its 186-482 part is similar to the 908-1254 part of Spike S; the 186-482 part of this human protein is included in the Pfam "Prominin" domain, Prominin/19-820. The other is the human prominin-2 protein (PROM2_HUMAN/NP_001159449) whose 174-473 part is similar to the 911-1254 part of Spike S. Note that when the "MSA generation iterations" parameter is set to 3 (default setting), similar results are obtained (thus, for example, the best result is the same in both cases).

For the given threshold of 0.90, one fly protein annotated with the Prominin domain of Pfam shares similarities with Spike S: the fly protein "Prominin-like protein" (PROML_DROME/NP_001261351.1, length = 1013) whose 235-534 part is similar to the 911-1254 part of Spike S; the 235-534 part of this fly protein is included in Pfam's "Prominin" domain, Prominin/76-881.

This suggests that part  908-1254 of Spike is a part of Prominim domain.

Note that for the probability threshold of 0.95, the human Prominin-1 shares a similarity with Spike glycoprotein of SARS-CoV. Its part 177-473 is similar to the part 890-1236 of Spike. The part 186-482 of this human protein is included in the Pfam domain “Prominin”, Prominin/19-820.

ORF3a share similarities with a “G-protein-coupled receptor” domain (robust similarity)

For the 0.80 probability threshold, only one match is identified: the human Lutropin-choriogonadotropic hormone (LSHR_HUMAN/NP_000224, length = 699). Its 537-693 part is similar to the 41-183 part of ORF3a. A large part of this 537-693 region is included in the Pfam domain "7 transmembrane receptor (rhodopsin family)", *i.e.* 7tm_1/376-623. Note that when the

"MSA generation iterations" parameter is set to 3 (default setting), no significant results are obtained (the probability of the best hit is 0.66).

For the given threshold of 0.80, no matches are detected with proteins belonging to the 4 proteomes tested.

However, for the probability threshold of 0.5, the search against the human proteome yields 28 hits including 26 G-protein coupled receptors:

- Lutropin-choriogonadotropic hormone receptor (LSHR_HUMAN/NP_000224, length = 699), proba = 80.39. Its part 537-693 is similar to the part 41-183 of ORF3a. A part of this 537-693 region is included in the Pfam domain “7 transmembrane receptor (rhodopsin family)”, *i.e.*, 7tm_1/376-623,

- “Probable G-protein coupled receptor 101” (GP101_HUMAN/NP_473362, length = 508), proba = 71.17. Its part 396-479 is similar to the part 66-138 of ORF3a. A large part of this 396-479 region is included in the Pfam domain “7 transmembrane receptor (rhodopsin family)”, *i.e.*, 7tm_1/48-454,

- Adhesion G-protein coupled receptor G5 (AGRG5_HUMAN/NP_001291305, length = 528), proba = 69.99. Its part 424-527 is similar to the part 45-137 of ORF3a. A large part of this 424-527 region is included in the Pfam domain “7 transmembrane receptor (Secretin family)”, *i.e.*, 7tm_2/245-499,

- Atypical chemokine receptor 2 (ACKR2_HUMAN/NP_001287, length = 384), proba = 63.95. Its part 227-337 is similar to the part 46-138 of ORF3a. A large part of this 227-337 region is included in the Pfam domain “7 transmembrane receptor (rhodopsin family)”, *i.e.*, 7tm_1/63-312,

- Muscarinic acetylcholine receptor M4 (ACM4_HUMAN/NP_000732, length = 479), proba = 63.79. Its part 397-477 is similar to the part 66-137 of ORF3a. A large part of this 397-477 region is included in the Pfam domain “7 transmembrane receptor (rhodopsin family)”, *i.e.*, 7tm_1/49-453,

- C-C chemokine receptor type 10 (CCR10_HUMAN/NP_057686, length = 362), proba = 63.45. Its part 224-335 is similar to the part 46-138 of ORF3a. A large part of this 224-335 region is included in the Pfam domain “7 transmembrane receptor (rhodopsin family)”, *i.e.*, 7tm_1/58-310,

- Alpha-2C adrenergic receptor (ADA2C_HUMAN/NP_000674, length = 462), proba = 62.39. Its part 379-462 is similar to the part 66-138 of ORF3a. A large part of this 379-462 region is included in the Pfam domain “7 transmembrane receptor (rhodopsin family)”, *i.e.*, 7tm_1/68-437,

- Melanocortin receptor 4 (MC4R_HUMAN/NP_005903, length = 332), proba = 59.41. Its part 233-327 is similar to the part 57-138 of ORF3a. A large part of this 233-327 region is included in the Pfam domain “7 transmembrane receptor (rhodopsin family)”, *i.e.*, 7tm_1/61-302,

- Melanopsin (OPN4_HUMAN/NP_150598, length = 478), proba = 58.76. Its part 246-375 is similar to the part 41-138 of ORF3a. A large part of this 246-375 region is included in the Pfam domain “7 transmembrane receptor (rhodopsin family)”, *i.e.*, 7tm_1/87-350,

- Adhesion G protein-coupled receptor E5 (AGRE5_HUMAN/NP_510966, length = 835), proba = 57.56. Its part 708-811 is similar to the part 46-138 of ORF3a. A large part of this 708-811 region is included in the Pfam domain “7 transmembrane receptor (Secretin family)”, *i.e.*, 7tm_2/546-781,

- Alpha-1D adrenergic receptor (ADA1D_HUMAN/NP_000669, length = 572), proba = 56.82. Its part 345-427 is similar to the part 66-138 of ORF3a. A large part of this 345-427 region is included in the Pfam domain “7 transmembrane receptor (rhodopsin family)”, *i.e.*, 7tm_1/113-402,

- Adhesion G protein-coupled receptor L4 (AGRL4_HUMAN/NP_071442, length = 690), proba = 55.99. Its part 576-688 is similar to the part 35-137 of ORF3a. A large part of this 576-688 region is included in the Pfam domain “7 transmembrane receptor (Secretin family)”, *i.e.*, 7tm_2/426-660,

- Adhesion G protein-coupled receptor L1 (AGRL1_HUMAN/NP_001008701, length = 1474), proba = 55.21. Its part 1009-1122 is similar to the part 35-138 of ORF3a. A large part of this 1009-1122 region is included in the Pfam domain “7 transmembrane receptor (Secretin family)”, *i.e.*, 7tm_2/859-1093,

- C-X-C chemokine receptor type 3 (CXCR3_HUMAN/NP_001495, length = 368), proba = 54.95. Its part 232-343 is similar to the part 46-138 of ORF3a. A large part of this 232-343 region is included in the Pfam domain “7 transmembrane receptor (rhodopsin family)”, *i.e.*, 7tm_1/70-318,

- D(1B) dopamine receptor (DRD5_HUMAN/NP_000789, length = 477), proba = 54.83. Its part 293-383 is similar to the part 66-138 of ORF3a. A large part of this 293-383 region is included in the Pfam domain “7 transmembrane receptor (rhodopsin family)”, *i.e.*, 7tm_1/757-359,

- Muscarinic acetylcholine receptor M1 (ACM1_HUMAN/NP_000729, length = 460), proba = 53.74. Its part 362-441 is similar to the part 66-136 of ORF3a. A large part of this 362-441 region is included in the Pfam 7tm_1 domain, *i.e.*, 7tm_1 /42-418.

- Thyrotropin-releasing hormone receptor (TRFR_HUMAN/NP_003292, length = 398), proba = 52.76. Its part 202-345 is similar to the part 41-138 of ORF3a. A large part of this 202-345 region is included in the Pfam 7tm_1 domain, *i.e.*, 7tm_1/42-320.

- Protein lunapark isoform 4 (LNP_HUMAN/NP_085153, length = 428), proba = 52.73. This protein does not belong to the GCPRs family.

- Atypical chemokine receptor 3 (ACKR3_HUMAN/NP_064707, length = 362), proba = 52.63. Its part 229-339 is similar to the part 46-137 of ORF3a. A large part of this 229-339 region is included in the Pfam 7tm_1 domain, *i.e.*, 7tm_1/62-315.

- Sphingosine 1-phosphate receptor 1 (S1PR1_HUMAN/NP_001307659, length = 382), proba = 52.39. Its part 244-336 is similar to the part 57-138 of ORF3a. A large part of this 244-336 region is included in the Pfam 7tm_1 domain, *i.e.*, 7tm_1/62-311.

- Probable G-protein coupled receptor 85 (GPR85_HUMAN/NP_001139737, length = 370), proba = 52.19. Its part 282-364 is similar to the part 66 -138 of ORF3a. A large part of this 282-364 region is included in the Pfam 7tm_1 domain, *i.e.*, 7tm_1/37-338.

- Alpha-2A adrenergic receptor (ADA2A_HUMAN/NP_000672, length = 465), proba = 52.13. Its part 377-464 is similar to the part 57-136 of ORF3a. A large part of this 377-464 region is included in the Pfam 7tm_1 domain, *i.e.*, 7tm_1/65-441.

- 5-hydroxytryptamine receptor 2B isoform 1 (5HT2B_HUMAN/NP_000858, length = 481), proba = 51.98. Its part 321-405 is similar to the part 66-138 of ORF3a. A large part of this 321-405 region is included in the Pfam 7tm_1 domain, *i.e.*, 7tm_1/71-380.

- 5-hydroxytryptamine receptor 6 (5HT6R_HUMAN/NP_000862, length = 440), proba = 51.95. Its part 193-345 is similar to the part 43-138 of ORF3a. A part of this 193-345 region is included in the Pfam 7tm_1 domain, *i.e.*, 7tm_1/44-320.

- Voltage-dependent calcium channel gamma-5 (CCG5_HUMAN/NP_665810, length = 275), proba = 51.94. This protein does not belong to the GCPRs family.

- C-X-C chemokine receptor type 4 isoform b (CXCR4_HUMAN/NP_003458, length = 352), proba = 51.33. Its part 216-327 is similar to the part 46-138 of ORF3a. A large part of this 216-327 region is included in the Pfam 7tm_1 domain, *i.e.*, 7tm_1/55-302.

- Muscarinic acetylcholine receptor M2 (ACM2_HUMAN/NP_001006627, length = 466), proba = 50.79. Its part 384-463 is similar to the part 66-136 of ORF3a. A large part of this 384-463 region is included in the Pfam 7tm_1 domain, *i.e.*, 7tm_1/40-440.

- Bombesin receptor subtype-3 (BRS3_HUMAN/NP_001718, length = 399), proba = 50.5. Its part 268-355 is similar to the part 66-138 of ORF3a. A large part of this 268-355 region is included in the Pfam 7tm_1 domain, *i.e.*, 7tm_1/64-328.

At the 0.50 probability level, two fly proteins share similarity with ORF3a. Both are G-protein coupled receptors:

- G-protein coupled receptor moody (MOODY_DROME/NP_001188534.1, length = 670), proba = 61.67. Its part 310-388 is similar to the part 66-138 of ORF3a. A large part of this 310-388 region is included in the Pfam 7tm_1 domain, *i.e.*, 7tm_1/53-363.

- Dopamine 1-like receptor 2, isoform B (DOPR2_DROME/NP_524548, length = 539), proba = 50.8. Its part 416-499 is similar to the part 66-138 of ORF3a. A large part of this 416-499 region is included in the Pfam 7tm_1 domain, *i.e.*, 7tm_1/125-474.
