## Supplemental file 3 for "Re-annotation of SARS-CoV-2 proteins using an HHpred-based approach opens new opportunities for a better understanding of this virus"

### Assessment of our results in the light of known weaknesses of HHpred

ORF3a:

We give below the UniProt queries used to perform our first proof:

Number of expressed human proteins (protein counts): **(proteome:UP000005640)** ; UP000005640 is the UniProt ID of Homo sapiens (Human): 80581

Number of human proteins labeled as "transmembrane": **(proteome:UP000005640) AND (keyword:KW-0812)** ; KW-0812 means “Transmembrane”: 13876.

Of the 28 proteins found by HHpred, 28 out of 28 are transmembrane proteins. The expected results are: (13876/80581)*32, or 4,821583252 and ((80581-13876)/80581)*28, or 23,178416748, which can be approximated by 5 and 23.

The contingency table is thus the following :

| | |
|---|---|
| 28 | 0 |
| 5 | 23 |

As more than 20% of the values are less than 5, the chi-square test cannot be applied. We therefore use a Fisher exact test instead of the chi-square test. The calculated Fisher p-value is 6.2059249716913E-11 (https://biostatgv.sentiweb.fr/?module=tests/fisher). Assuming that the null hypothesis is rejected for a contingency table that is assigned a p-value of 5% or less by the Fisher test, we conclude that the results found by HHpred are not randomly drawn from the UniProt human proteome.

Second proof.

We give below the UniProt queries used to perform our second proof:

Number of human proteins annotated with the 7tm_1: **(proteome:UP000005640) AND (xref:pfam-PF00001)** ; PF00001 corresponds to the Pfam 7tm_1 domain: 412

Number of human proteins annotated with the 7tm_2: **(proteome:UP000005640) AND (xref:pfam-PF00002)** ; PF00002 corresponds to the Pfam domain 7tm_2: 128

Number of human proteins annotated with both the 7tm_1 and 7tm_2 domains of Pfam: **(proteome:UP000005640) AND (xref:pfam-PF00001) AND (xref:pfam-PF00002)**: 0

Of the 28 transmembrane proteins found by HHpred, 26 are 7tm_1 or 7tm_2, the other two are transmembrane proteins that do not belong to these two classes. The expected results are: (540/13876)*32, *i.e.* 1,089651196 and ((13876-540)/ 13876)*32, *i.e.* 26,910348804, which can be approximated by 1 and 27.

The contingency table is therefore as follows:

| | |
|---|---|
| 26 | 2 |

| | |
|---|---|
| 1 | 27 |

As more than 20% of the values are less than 5, the chi-square test cannot be applied. We therefore use a Fisher exact test instead of the chi-square test. The computed Fisher’s p-value is 2.8739559680731E-12 (https://biostatgv.sentiweb.fr/?module=tests/fisher). Assuming that the null hypothesis is rejected for a table assigned a p-value of 5% or less by Fisher's test, we conclude that the results found by HHpred are not randomly drawn from the class of the transmembrane protein family comprising the 7tm_1 or 7tm_2 proteins compared to the other classes.
