## Supplemental file 5 for "Re-annotation of SARS-CoV-2 proteins using an HHpred-based approach opens new opportunities for a better understanding of this virus": README

File hhpred_7375456.hhr corresponds to the query using the entire sequence of "Lutropin".
File hhpred_7375456.hhr corresponds to the query using the region of "Lutropin" that shares homology with a portion of SARS-CoV-2 ORF3a.
